# Conditional deletion of *Tmem63b* does not impact mouse voiding behavior

**DOI:** 10.1101/2025.04.07.647591

**Authors:** Marianela G. Dalghi, Wily G. Ruiz, Dennis R. Clayton, Tanmay Parakala-Jain, Marcelo D. Carattino, Yun Stone Shi, Gerard Apodaca

## Abstract

The proper function of the lower urinary tract depends on its ability to sense and react to mechanical forces as urine is produced, transported, stored, and eliminated; however, our current understanding of the mechanosensors involved in these events is limited. TMEM63 ion channels are reported to function as mechanosensors/osmosensors in other organs, and our studies revealed that the primary site of *Tmem63a* and *Tmem63b* gene expression and TMEM63B protein expression in the mouse bladder wall was the urothelium. Despite this localization, voiding behavior in conditional urothelial *Tmem63b* knockout mice, assessed using a video-monitored void-spot screening assay, was not significantly different from control mice, even when the urothelium was stressed by exposure to cyclophosphamide. We further observed that dorsal root ganglia sensory neurons, including those innervating the bladder, were also sites of *Tmem63a*, *Tmem63b*, and TMEM63B expression. Again, voiding behavior was not impacted in conditional sensory neuron *Tmem63b* knockout mice, treated or not with cyclophosphamide. Our studies reveal that the urothelium and dorsal root ganglia are sites of *Tmem63a*, *Tmem63b*, and TMEM63B expression, but deletion of *Tmem63b* alone in these tissues does not result in a demonstrable voiding phenotype.

## INTRODUCTION

The normal function of the lower urinary tract depends on mechanisms to sense and respond to mechanical forces as urine traverses the renal pelvis, ureters, bladder, and urethra. These forces include: shear stress as newly formed urine is propelled across the mucosal surfaces of the ureter; tensile forces that arise in the lining urothelium and subjacent tissues as the bladder fills with urine; and compressive forces that are generated as the detrusor contracts and the mucosal surface of the bladder refolds in response to voiding. Perturbations in these events may underlie or contribute to several lower urinary tract pathologies including neurogenic bladder, partial outlet obstruction, and conditions such as overactive bladder and underactive bladder. Although all tissues in the bladder wall are reported to be mechanosensitive [1–9], much research has focused on the interface between the urothelium and subjacent nerve processes. A current hypothesis is that bladder filling triggers the serosal release of urothelial mediators (e.g., ATP, adenosine, NO, acetylcholine), which upon binding to subjacent sensory neuron processes relay information about the filling status of the bladder to the central nervous system [10–13]. Yet, we have an incomplete understanding of the molecules that lower urinary tract tissues use to sense mechanical stimuli, the mechanotransduction pathways involved, or the contributions they make to normal and abnormal physiology. Thus, an important research goal is to identify lower urinary tract tissue-associated mechanosensors and assess their impact on voiding function.

Mechanosensors are biological force transducers that directly sense and respond to mechanical stimuli by undergoing conformational changes, ultimately triggering downstream mechanotransduction cascades [14–16]. Numerous classes of mechanosensors have been described including apical specializations (e.g., cilia), G-protein coupled receptors, elements of the cytoskeleton, proteins associated with cell-cell and cell-matrix junctions, and ion channels [14, 17–26]. Examples of the latter include the PIEZO family channels PIEZO1 and PIEZO2, members of the two-pore K^+^ channel family including KCNK2 (TREK-1), KCNK10 (TREK-2), KCNK4 (TRAAK), and orthologs of the plant osmosensitive OSCA1 channels identified in the mouse by Zhao *et al.* including TMEM63A and TMEM63B [27, 28]. The latter are classified as high threshold, low conductance non-selective cation channels that are mechano- and osmo-sensitive [29–31].

TMEM63 family members are implicated in a variety of cellular functions and disease processes. In *Drosophila melanogaster*, a single TMEM63 ortholog is required for detection of humidity by Or42b neurons and detection of food texture by the md-L neurons that innervate sensory sensilla [32, 33]. Also in *Drosophila,* as well as in the mouse neuroblastoma Neuro-2A cell line, orthologs of TMEM63A reportedly function as lysosomal mechanosensors [34]. *Tmem63a* may modulate chronic post-amputation pain [35], and mutations in *TMEM63A* are associated with transient hypomyelination during infancy and hypomyelinating leukodystrophy [36–39]. TMEM63A in conjunction with TMEM63B function as mechanosensors in stretch-induced surfactant and ATP release by alveolar pneumocytes [40]. TMEM63B-dependent osmosensation is required for vertebrate hearing [29] and is also integral to thirst perception and detection of hyperosmolarity in the subfornical organ of the brain [41], the secretion of insulin [42], and release of thyroid hormone [43]. Gain-of-function variants of *TMEM63B* are associated with early-onset epilepsy, progressive brain damage, and hematological disorders [44]. An additional isoform, TMEM63C, has been described and is implicated in hereditary spastic paraplegia as well as defects in kidney podocyte function [45, 46]; however, it does not function as a mechanosensitive channel when reconstituted in liposomes [30]. What TMEM63C is doing in these contexts is unknown and awaits further exploration.

Our studies reveal that the urothelium and dorsal root ganglia (DRG; comprised of sensory neurons and accessory cells) are sites of TMEM6B/*Tmem63b* expression. To test the hypothesis that TMEM63B is integral to urothelial mechanotransduction, we generated conditional urothelial *Tmem63b* knockout (KO) mice and assessed their voiding behavior using video-monitored void spot assays. However, these mice had no observable voiding phenotype.

Likewise, conditional sensory nerve *Tmem63b* KO mice exhibited no voiding defects. We discuss the implications of these findings, including the possibility that *Tmem63a,* which is also expressed by the urothelium and sensory neurons, compensates for loss of *Tmem63b* expression in these tissues.

## MATERIALS and METHODS

### Animals and PCR genotyping

*Tmem63b^HA-fl/HA-fl^* mice, previously described by Du et al. [29], were generated by using CRISPR to insert nucleotides that encode an N-terminal HA tag (and 3X FLAG tag) after the 20-amino acid signal sequence. LoxP sites were also inserted into introns 1 and 4. The mice were backcrossed and maintained on a C57BL/6 background. Wild-type C57BL/6J mice, which we refer to as *Tmem63b^+/+^*mice in this manuscript, were obtained from the Jackson Laboratory (strain 000664; Bar Harbor, ME) and used as controls in some of our studies. *Avil^Cre+/–^* mice and *Upk2^Cre+/–^* mice were obtained from Jackson Laboratory (strains 032536 and 029281, respectively) [47, 48]. Conditional urothelial *Tmem63b* KO mice were generated by crossing *Tmem63b^HA-fl/HA-fl^* mice with *Tmem63b^HA-fl/HA-fl^*;*Upk2^Cre+/–^*mice. Conditional sensory neuron *Tmem63b* KO mice were generated by crossing *Tmem63b^HA-fl/HA-fl^* mice with *Tmem63b^HA-fl/HA-fl^*;*Avil^Cre+/–^* mice. Expression of *Upk2* in all cell layers of the mouse bladder urothelium was confirmed by mating *Upk2^Cre+/–^* mice with Ai9 reporter mice (Jackson Laboratory strain 007909), which express CAG promoter-driven tandem-dimer-Tomato red fluorescent protein after Cre- mediated removal of a stop codon. Mice were housed in standard solid-bottom caging with up to five females/cage and up to four non-breeding males/cage. Animals were provided with paper blocks to generate bedding/nesting sites. Pregnant females were housed separately, as were male breeders. Mice were housed under a 12-hour day/night cycle and were fed standard mouse chow (Labdiet 5P76, irradiated; Purina, Wayne County, IN) and given water *ad libitum*. Mice were harem bred, and breeding males were housed one/cage.

Genotyping was performed on tail snips collected from 21- to 25-day old pups. Tail DNA was extracted using the HotSHOT NaOH method [49] and PCR reactions were performed using the KAPA2G Fast Hot Start Ready Mix with dye (cat# 07961278001, Roche Diagnostics, Indianopolis, IN). *Tmem63b^HA-fl/HA-fl^* was detected using the following primers: forward – TCA ACA GCA GCA ACC CGA AG; reverse – CAC ATG AAG TCC AGA GCC AG [50]. The following PCR cycling conditions were employed: initial denaturation was 95°C for 5 min, followed by 35 cycles of 95°C for 30s, 56°C for 30s, 72°C for 20s, and a final step of 72°C for 7 min. *Avil^Cre^*and *Upk2^Cre^* expression was confirmed using the primers and PCR reaction protocols described previously and by Jackson Laboratories [2, 29]. Experiments were performed with females and males between 9- and 24-weeks old. Animal studies were performed in accordance with relevant guidelines/regulations of the Public Health Service Policy on Humane Care and Use of Laboratory Animals and the Animal Welfare Act, and under the approval of the University of Pittsburgh Institutional Animal Care and Use Committee (protocol number 24044849). Mice were euthanized by CO_2_ asphyxiation, followed by thoracotomy as a secondary method.

### Reagents, antibodies, and fluorescent probes

Unless specified otherwise, all chemicals were obtained from Sigma-Aldrich (St Louis, MO). Rabbit monoclonal antibody (C29F4) to the HA epitope was purchased from Cell Signaling Technology (Danvers, MA; catalog number 3724S). Rabbit anti-cholera toxin B subunit antibody was from Novus (Centennial, CO; catalog number NB100-63607). The following secondary antibodies were purchased from Jackson ImmunoResearch Laboratories (West Grove, PA): donkey anti-rabbit-Alexa Fluor 488 (catalog number 711-545-152), goat anti-rabbit-Alexa Fluor 488 Fab fragments (catalog number 111-547-003), and goat anti-rabbit-CY3 (catalog number 111-165-144). Alexa Fluor 488- or rhodamine-labeled phalloidin (catalog numbers A12379 and R415, respectively) and DAPI (catalog number 62248) were obtained from ThermoFisher Scientific (Grand Island, NY).

### PCR analysis

Freshly excised mouse bladders were rinsed with Kreb’s buffer (110mM NaCl, 25mM NaHCO_3_, 5.8mM KCl, 1.2mM MgSO_4_, 4.8mM KH_2_PO_4_, 11mM glucose, 2mM CaCl_2_, gassed with 5% v/v CO_2_). Fine forceps were used to invert the bladder onto the pointed end of a yellow tip, trimmed by 5mm with a scalpel, that was positioned next to the dome of the bladder. The inverted bladder was placed in 150µl of lysis/binding buffer (RNAqueous-4PCR Kit; Invitrogen, Waltham, MA) for 30s and total RNA was isolated per the manufacturer’s protocol. An AccuSript PfuUltra II RT-PCR kit (Agilent, Santa Clara, CA) was used to generate cDNAs using random primers and following the manufacturer’s protocol. The following primers were used to identify gene expression by PCR:

***Tmem63a*-FWR** TGAAAACGAGCTGGGATGCT

***Tmem63a*-REV** CCTTTCTGGCTTCTCTGGGG

***Tmem63b*-FWR** CTGCGCCTAGGGGAGGAT

***Tmem63b*-REV** CGGAGGTGAGACGCTCATAC

***Tmem63c*-FWR** GAGAGCAAGTTCCTGTGGCT

***Tmem63c*-REV** GCTCCGCATCTACCTCCTTG

PCR reactions were performed using the KAPA HiFi polymerase kit (Roche Sequencing Solutions, Pleasanton, CA) and a T100 Thermal Cycler (BioRad, Hercules, CA) using the following protocol: initial denaturation at 94°C for 2min, followed by 39 cycles of 94°C for 20s, 60°C for 15s, 72°C for 15s, and a final step of 72°C for 2min prior to holding at 4°C. Amplicons were resolved using 2% w/v agarose gels.

### Fresh-frozen sample preparation

Mice were euthanized by CO_2_ asphyxiation, their abdominal hair moistened using 70% v/v ethanol, and a caudal midline abdominal incision made using a sterile scalpel. The bladder was exteriorized and released from the body by cutting through its neck region just above the urethral sphincter using sharp tissue scissors. The bladder was rinsed in Kreb’s buffer, plunged in Optimal Cutting Temperature (OCT) solution (Tissue-Tek, Sakura Finetek, Torrance, CA) to remove excess buffer, and then placed dome-side down in cryomolds (15 x 15 x 5 mm; Fisher Scientific) filled with OCT. The samples were flash frozen by placing the cryomold on metal plate cooled with liquid nitrogen and stored at -80°C in sealed plastic bags. Cryosections were cut using a Leica Microsystems CM1950 cryostat (Buffalo Grove, IL; 8-12 µm sections; -20°C chamber and -18°C knife temperatures), collected on Superfrost Plus glass slides (ThermoFisher Scientific, Pittsburgh, PA), and held within the cryochamber at -20°C for 30 min (to promote adherence of the section to the slide) prior to fixation/staining. Alternatively, slides were stored at -80°C.

### Detection of gene expression by fluorescent in situ hydridization (RNAscope) and by BaseScope

RNAscope probes and associated multiplex RNAscope and Basescope kits were obtained from Advanced Cell Diagnostics (ACD; a Biotechne brand; Newark, CA). The following ACD probes were used for RNAScope analysis: *Tmem63a*-channel 1 (catalog number 431521), *Tmem63b*- channel 2 (catalog number 431531-C2), *Krt8*-channel 3 (catalog number 424521-C3), and *<u>B</u>acillus subtilis dapB* (catalog number 320871). *Krt8,* which is expressed by all urothelial cell layers, served as a positive control and *dapB* as a negative one. The ACD protocol for fresh-frozen tissue sections was employed with the following changes. The frozen sections, prepared as described above, were fixed by adding cold (4°C), freshly prepared neutral buffered formalin (4% w/v paraformaldehyde dissolved in 29.04 mM NaH_2_PO_4_•H_2_O and 45.8 mM Na_2_HPO_4_, pH 7.4) and incubation for 15 min at room temperature. Protease IV treatment for 5 min at room temperature was employed. Opal reagent packs (Akoya Biosciences; Marlboro, MA) were used to fluorescently label RNAs: The Opal520 reagent pack for *Tmem63a*, Opal570 reagent pack for *Tmem63b*, and Opal690 reagent pack for *Krt8*. The Opal reagents were dissolved in 75µl of the provided DMSO, stored at 4°C, and used at a 1:1000 dilution during the RNAScope labeling protocol. Slides incubated with positive or negative control probes were run side-by-side and treated identically. Labeled tissue was mounted using ProLong Gold antifade mountant (ThermoFisher), cured overnight at room temperature in the dark, and the slides subsequently stored at 4° C prior to viewing and image acquisition.

To detect *Upk2^Cre^*-mediated recombination of *Tmem63b^HA-fl/HA-fl^*, a custom BaseScope probe called BA-Mm-Tmem63b-3zz-st-C1 was generated (catalog number 1129311-C). This probe was designed to detect the presence of a 218bp fragment encoding exons 2-4 prior to Cre-mediated excision. The ACD BaseScope protocol was followed using fresh frozen tissue that was fixed and protease treated as described above. Positive and negative BaseScope control probes (catalog number 322976) were run concurrently. Images were collected using an HC PLAN APO 10X objective (N.A. 0.40) or an HCX PLAN APO 40X oil objective (N.A. 1.25) attached to a DM6000B widefield microscope (Leica Microsystems, Buffalo Grove, IL) outfitted with a Gryphax Prokyon digital camera (Jenoptik, Jupiter, FL), and using an Apple (Cupertino, CA) iMac computer running Gryphax software (Jenoptik). Images (1920 x 1200 pixels) were saved in TIFF file format. To quantify *Upk2-Cre*-dependent recombination, we opened random images of BaseScope-labeled tissues in Fiji (ImageJ2) and then used the freehand tool to mark the region of interest (ROI; i.e., urothelium). The area of the ROI was recorded, and the Cell Counter plugin was used to count the number of positive *Tmem63b* Basescope dots in this ROI. The number of dots per mm^2^ of urothelial area were calculated and recorded in a Microsoft (Redmond, WA) Excel spreadsheet. For figures, images were contrast corrected in Adobe (San Jose, CA) Photoshop 2025 and composite images generated in Adobe Illustrator 2025.

### Confocal microscopy and image processing

Images were captured by confocal microscopy using either a Leica HCX PL APO 20X, 0.75 NA dry objective or a Leica HCX PL APO CS 40X, 1.25 NA oil objective (attached to a Leica DMI8 microscope) and the appropriate laser lines of a Leica Microsystems SP8 Stellaris confocal system outfitted with a 405-laser diode and a white-light laser. The signal from the Power HyD detectors was optimized using the Q-LUT option, and 8-bit images collected at 400-600 Hz using 3-line averages. Crosstalk between channels was prevented by use of spectral detection coupled with sequential scanning. Stacks of images (1024 x 1024, 8-bit) were collected using system-optimized parameters for the Z-axis. Images were processed using the 3D visualization option in Bitplane Imaris (Boston, MA) and exported as TIFF files. If necessary, the contrast of the images was corrected in Photoshop CC2025, and composite images prepared in Adobe Illustrator CC2025.

### Labeling afferent neurons with cholera toxin (Ctx) and isolation of DRGs

*Tmem63b^HA-fl/HA-^*^fl^ mice were anesthetized using isoflurane (3% v/v), the abdomen sterilized with povidone iodine, a caudal midline incision was made using a scalpel, and the bladder was exteriorized. Ctx beta subunit (MilliporeSigma; Burlington, MA; catalog number C9903), dissolved at 0.5% w/v in sterile 0.9% w/v saline, was injected into the bladder wall at four sites (2 µl/injection site) using a Hamilton Company (Reno, NV) model 701 gastight microliter syringe outfitted with a 33g needle (Hamilton catalog number 7803-15). The abdominal incision was closed in layers using 5.0 polydioxanone absorbable monofilament surgical sutures (AD Surgical, Sunnyvale, CA, USA). Ketoprofen (5 mg/kg; Zoetis, Parsippany, NJ, USA) was administered subcutaneously to alleviate pain and ampicillin (100 mg/kg; Eugia, Hightstown, NJ, USA) was given to prevent infections. DRG from L4-L5 or those at L6-S2 were harvested 7 days later using our previously described protocols [51, 52]. The DRGs from three mice were pooled, frozen in OCT compound, and sectioned as described above for bladder preparations.

### Detection of HA-TMEM63B and Ctx in tissues by immunofluorescence

Detection of HA-TMEM63B by immunofluorescence was performed as described previously [50]. Frozen sections of bladder and DRG (prepared as described above) were fixed by adding cold (4°C), freshly prepared neutral buffered formalin and incubation for 10 min at room temperature. The tissue was rinsed with phosphate buffered saline (PBS; 137mM NaCl, 2.7mM KCl, 8.1mM Na_2_HPO_4_, and 1.5 mM KH_2_PO_4_, pH 7.4) and unreacted fixative was quenched by incubating the tissue slices for 10 min at room temperature with Quench Buffer (75 mM NH_4_Cl and 20 mM glycine, pH 8.0 dissolved in PBS, containing 0.1% v/v Triton X-100). The tissue was then rinsed 3 times with PBS followed by incubation in Block Solution (PBS containing 0.6% v/v fish skin gelatin and 0.05% w/v saponin) containing 10% v/v donkey serum for 60 min at room temperature in a humid chamber. The Block Solution was aspirated and replaced with primary antibodies diluted in Block Solution and incubated for 1 h at room temperature (or overnight at 4° C) in a humid chamber. The slides were washed 3-times quickly and 3 times for 3 min with Block Solution, and then incubated with fluorophore-labeled secondary antibodies, diluted in Block Solution, for 1 h at room temperature. DAPI (1:1000) and rhodamine-phalloidin (1:200) were included during the secondary antibody incubation. The labeled tissues were then rinsed 3-times quickly and 3 times for 5 min with Block Solution, rinsed with PBS, and then postfixed in neutral buffered formalin for 5-10 min at RT. The slides were rinsed with PBS, excess liquid aspirated, and a drop of SlowFade Diamond Antifade was placed on the tissue. Borosilicate coverslips (#1.5, 0.17 mm thickness, 24 x 50 mm; ThermoFisher) were placed above the drop of mounting medium, excess mounting medium was removed by aspiration, the edges of the coverslip were sealed with clear nail polish, and after the nail polish dried the slides were stored at -20 °C until image acquisition was performed. Control incubations lacked primary antibodies or secondary antibodies.

When simultaneously immunolocalizing Ctx and HA-TMEM63B, we used two rabbit primary antibodies. Following tissue fixation in neutral buffered formalin, the tissue was rinsed with PBS, then with Quench Solution, and subsequently incubated in Block Solution containing 5% v/v goat serum for 1 h at room temperature. The tissue was washed 3 times 3 min with Block Solution and then incubated with first primary antibody, rabbit anti-HA (diluted 1:100 in Block Solution), for 1 h at room temperature. The tissue was then washed 3 times 3 min with Block Solution and incubated for 1 h at room temperature with the first secondary antibody (goat anti-rabbit-Alexa Fluor 488 Fab fragments) diluted 1:500 in Block Solution. The tissue was then washed for 3 times 5 min with PBS. The antibodies were fixed using neutral buffered formalin for 10 min at room temperature, rinsed with PBS and then Quench buffer. The tissue was incubated with Block Solution for 5 min and then reacted with the second primary antibody (rabbit anti-Ctx B subunit) diluted 1:500 in Block Solution for 60 min. The tissue was washed 3 times 5 min with Block Solution and then incubated with the second secondary antibody (CY3- conjugated goat-anti-rabbit antibody; 1:3000 dilution) for 60 min at room temperature. The samples were then washed 3 times 5 min with Block Solution, rinsed with PBS, and then post-fixed and mounted as described above. Control incubations included the following: (1) omission of the first primary antibody; (2) omission of the second primary antibody; (3) omission of the first primary antibody, the first secondary antibody, and the second primary antibody; (4) omission of the first primary antibody, the second primary antibody, and the second secondary antibody.

### Video-monitored void spot analysis

To assess voiding behavior in awake, freely moving mice, we used previously described custom-built void-spot chambers outfitted with video monitoring [2, 53]. These chambers included an upper compartment that housed an individual mouse with continuous access to food and water (in the form of Hydrogel; ClearH2O, Westbrook, ME) and an igloo-shaped sleeping chamber. The floor of the upper chamber was lined with chromatography/blotting paper and illuminated from below by UV lights housed in a lower chamber. The animals and paper were monitored by wide-angle “webcam” cameras positioned above the upper chamber and another mounted at the base of the lower compartment. The incorporation of real-time video monitoring allowed us to follow mouse activity over extended periods of time while overcoming the difficulties of distinguishing overlapping voiding spots, a shortfall of standard void-spot assays [54]. The mice were routinely housed in a facility with 12-hour light-dark cycles, with 7:00AM being zeitgeber time (ZT)=0 (start of light cycle). To analyze their voiding behavior in the dark phase, the mice were placed in the upper chamber between 17:00-18:00 h (ZT10-11), and analysis of void spots was performed from 00:00-6:00 (ZT=17-23). The extended period of acclimatization reflected access limitations to the facility after 7:00 PM. For analysis during the light phase, the mice were placed in the chamber between 10:00-11:00 (ZT=3-4), allowed to acclimatize for one hour, and analysis performed during the subsequent 6-h time window. Video was captured at 1 frame per second with a 1920 × 1080 pixel resolution using an Apple M3 Mac mini-computer running SecuritySpy software (BenSoftware.com). The movies were saved in .m4v format and viewed on an Apple Intel i9 iMac computer using Quicktime (Apple) software. Calibration curves, made by spotting mouse urine (2 µl – 750 µl) on the paper, were used to calculate the volume/void spot. As previously [2, 53], spots were categorized as primary void spots (PVS; those ≥ 20 µl) or secondary void spots (SVS; < 20 µl). The parameters measured in our analysis were number of PVS, average PVS volume per void, total PVS volume, number of SVS, and total SVS volume. We also calculated a ratio of total PVS volume ÷ (total PVS volume + total SVS volume). In untreated animals, the majority of void spots were PVS and the score (referred to as tPVS/(tPVS+tSVS) in the figures) was ∼ 1.0. However, in animals treated with cyclophosphamide (CYP), the inflammation resulted in an overactive bladder phenotype in which increasing amounts of the total volume were contributed by SVS, reducing the overall ratio. Animals that chewed or damaged the paper prior to or during the time window of analysis were excluded from the study.

### Cyclophosphamide treatment and analysis

An acute (single dose) cyclophosphamide (CYP) model was employed. A 25 mg/ml stock of CYP, dissolved in sterile 0.9% w/v saline, was prepared daily and filtered through a sterile 0.22 μm syringe filter. In unpaired experiments, mice were injected once intraperitoneally with CYP (150 mg/kg; i.e., 6 µl of 25 mg/ml stock injected per g mouse weight) or with vehicle (0.9% w/v saline) at 10:00-11:00 (ZT=3-4) and void-spot analysis performed in the subsequent 0:00-6:00 (ZT=17-23) time frame. The effects of CYP (i.e., increased urination) are observable within one hour of injection. In paired experiments, mice were injected with saline at 10:00-11:00 (ZT=3-4) and void spot analysis performed in the subsequent 0:00-6:00 (ZT=17-23) time window. Next, the animals were injected with CYP at 10:00-11:00 (ZT=3-4) and void-spot analysis performed from 0:00-6:00 (ZT=17-23).

### Statistical analysis

Data are expressed as mean ± SEM (*n*), where *n* equals data from an individual mouse. Parametric or nonparametric tests were employed as appropriate. CYP experiments were analyzed using 2-way ANOVA of log transformed data where Y=log(Y+1). Values of *p* ≤ 0.05 were considered statistically significant. Statistical comparisons were performed using GraphPad Prism 10 (GraphPad Software, San Diego, CA, USA).

## RESULTS

### Expression of Tmem63a and Tmem63b/TMEM63B in the mouse bladder

An important site of mechanotransduction in the bladder wall is the urothelium [1, 13]. We used RT-PCR to demonstrate that *Tmem63a* and *Tmem63b* were both expressed in the bladder and specifically in the urothelium (Fig. 1A). In contrast, we could not detect expression of *Tmem63c* in these tissues (Fig. 1A). These observations fit well with transcriptomic studies that report expression of *Tmem63a* and *Tmem63b* in the cells that populate the bladder, but not *Tmem63c* [55]. Fluorescent *in situ* hybridization (FISH) revealed that *Tmem63a* expression was concentrated in the urothelium, with scattered expression detected in the lamina propria and the muscularis externa (Fig. 1B-C). *Tmem63b* expression was also concentrated in the urothelium, but with less signal apparent in suburothelial tissues (Fig. 1B-C). Limited reaction product was detected in samples incubated with the *B. subtilis dapB* triple-negative control probe (Fig. 1B-C). To confirm TMEM63B protein expression in the urothelium, we used a previously published reporter mouse (*Tmem63b^HA-fl/HA-fl^*), which expresses an HA tagged version of TMEM63B (which we refer to as HA-TMEM63B) in place of the endogenous protein [29]. Consistent with our FISH analysis, we observed that HA-TMEM63B was highly enriched in the urothelium and not the other tissues of the bladder wall (Fig. 2A). Within the urothelium, the intensity of staining was polarized, with the greatest signal in the basal cell layer and decreasing amounts of signal as one progressed from intermediate cell layer to the outermost umbrella cell layer (see right-most panels of Fig. 2D). As a control for these studies, we demonstrated that HA-TMEM63B was not detected in the urothelium of wild-type (*Tmem63b^+/+^*) mice processed identically (Fig. 2A). A reporter mouse expressing a tagged version of *Tmem63a* was not available, so similar studies could not be performed for this gene product.

**Fig. 1.**
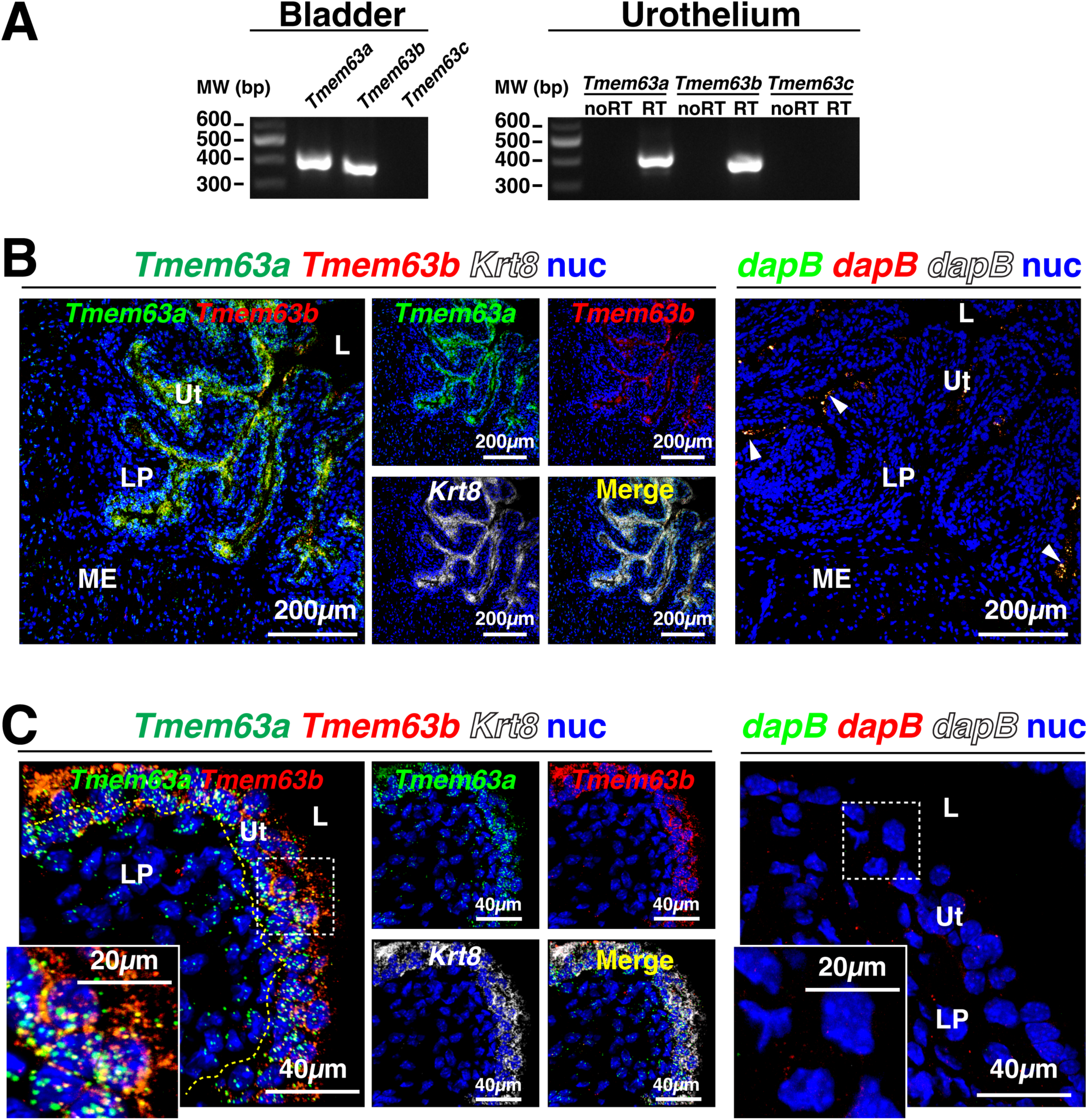
Expression of *Tmem63a* and *Tmem63b* in the bladder urothelium. **(A)** mRNA was isolated from whole bladder (left-most panel) or urothelium (right-most panel), reverse transcribed, and PCR used to detect the presence of the indicated gene product. **(B-C)** Expression and localization of message for the indicated gene as assessed by fluorescent *in-situ* hybridization. The *Krt8* probe, which labels all three layers of the urothelium, served as a positive control. The *dapB* probe serves as a negative control. Lower magnification overviews are provided in B and higher magnification views are included in C. The arrowheads in the right-most panel of B indicate umbrella cell-associated lysosomes, which variably exhibit autofluorescence. The boxed regions in C are magnified in the insets. *Legend:* L, lumen; LP, lamina propria; ME, muscularis externa; Ut, urothelium.

**Fig. 2.**
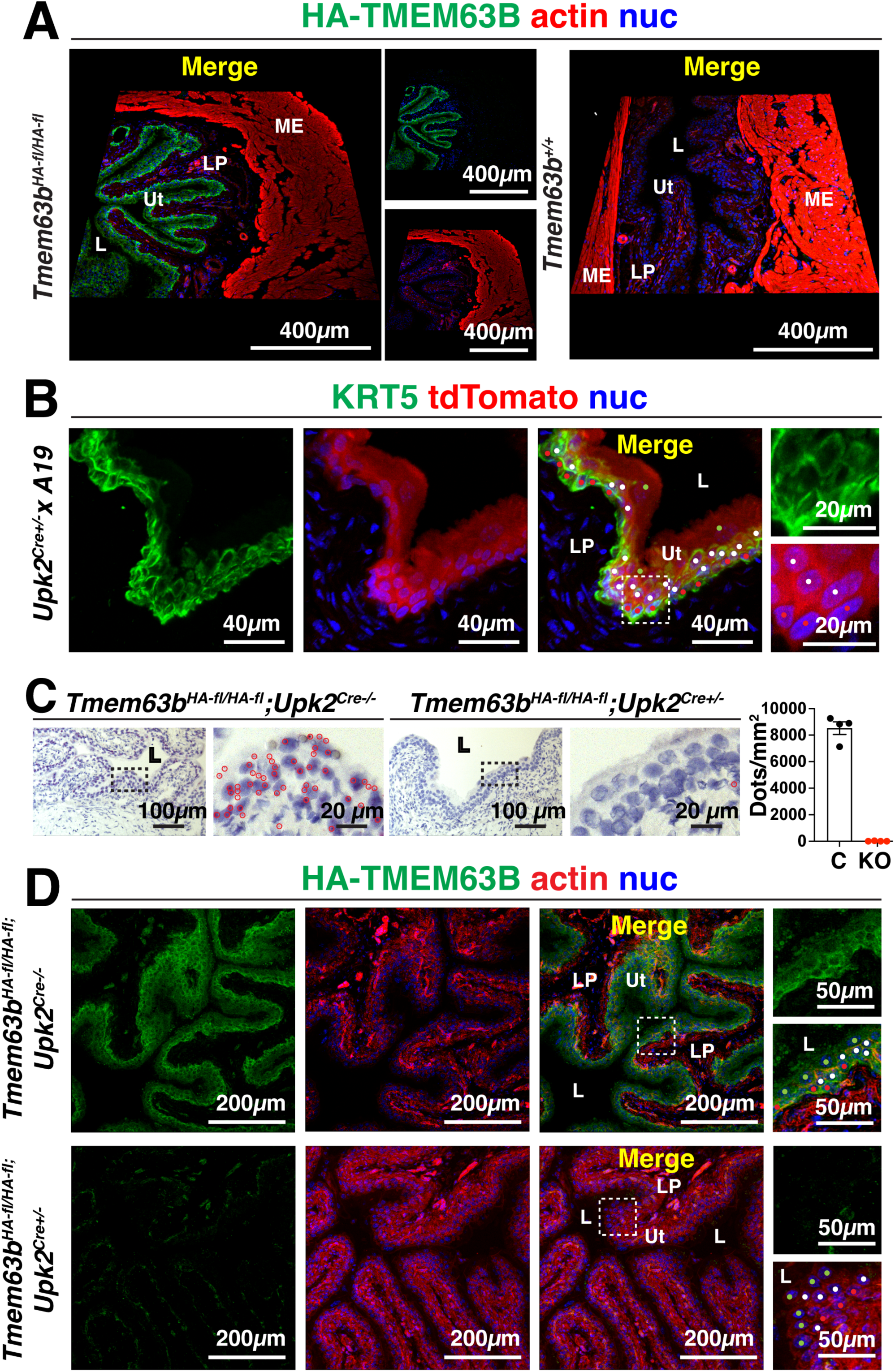
Expression of HA-TMEM63B/*Tmem63b* in conditional urothelial *Tmem63b* KO mouse bladders. **(A)** Localization of HA-TMEM63B in the bladders of *Tmem63b^HA-fl/HA-fl^* and wild-type *Tmem63B^+/+^* mice. **(B)** *Upk2^Cre+/-^*mice were crossed with Ai9 reporter mice and expression of tandem dimer (td)Tomato in the KRT5-labeled urothelial cells was assessed by immunofluorescence. The boxed region is magnified in the panels to the right. **(C)** Use of BaseScope to assess *Upk2-*Cre-mediated recombination of *Tmem63b^fl-HA/fl-HA^*exons 2-4. Reaction products (dots), indicating expression of *Tmem63b* exons 2-4, are marked with red circles. In the panel to the right, the number of dots per mm^2^ of urothelium was quantified in *Tmem63b^HA-fl/HA-fl^*;*Upk2^Cre-/-^*(control; C) and *Tmem63b^HA-fl/HA-fl^*;*Upk2^Cre+/-^*(conditional knockout; KO) mouse bladders. Data are mean ± SEM (n=4). **D.** HA-TMEM63B expression in *_Tmem63b_HA-fl/HA-fl*_;_ *Upk2^Cre-/-^* and *Tmem63b^HA-fl/HA-fl^*;*Upk2^Cre+/-^* mice. In panels B and D, red dots indicate basal cells, white dots indicate intermediate cells, and green dots indicate umbrella cells. *Legend:* L, lumen; LP, lamina propria; Ut, urothelium.

### Generation of conditional urothelial specific Tmem63b KO mice

Given the preponderance of *Tmem63b/*TMEM63B expression in the urothelium, combined with the availability of *Tmem63b^HA-fl/HA-^*^fl^ mice, we next set out to make conditional urothelial *Tmem63b* KO mice. Within the bladder, *Upk2^Cre^* mice are reported to drive selective expression of *Cre* in the mouse urothelium [47, 56], and we confirmed that that *Upk2^Cre^* mice employed in our studies drive Cre expression in all three strata of the urothelium (i.e., umbrella cell layer, intermediate cell layers, and basal cell layer) (Fig. 2B). To confirm knockout of *Tmem63b* in the urothelium, we used an *in situ* hybridization approach (BaseScope), which confirmed almost complete Cre-mediated excision of exons 2-4 in the urothelium of conditional urothelial *Tmem63b* KO mice (*Tmem63b^HA-fl/HA-fl^*;*Upk2^Cre+/-^*) versus control ones (*Tmem63b^HA-fl/HA-fl^*;*Upk2^Cre-/-^*)(Fig. 2C). Furthermore, we observed a loss of urothelial HA-TMEM63B protein expression in control versus conditional urothelial KO mice (Fig. 2D).

### Voiding behavior is not affected in conditional urothelial Tmem63b KO mice

A useful screening tool for detecting defects in lower urinary tract function, including alterations in mechanotransduction, is the void spot assay [2, 3, 57–59]. We recently developed a video-monitored version of this approach and used it to assess voiding behavior in male and female control and conditional urothelial *Tmem63b* KO mice during their active (dark) and resting (light) phases [2, 53]. As described previously, we binned voids into primary voids spots (PVS; defined as ≥ 20 µl) or secondary void spots (SVS; defined as < 20 µl) [2, 53]. The former are associated with normal voiding behavior, while the latter are somewhat more prevalent in dominant males and are significantly increased in animals (regardless of sex) with an overactive bladder phenotype [54, 60–62]. However, we did not observe a significant difference in PVS parameters (PVS number, average PVS volume, and total PVS volume), SVS parameters (SVS number and total SVS volume), or fraction of total urinary output contributed by PVS, i.e. the tPVS/(tPVS+tSVS) ratio, when comparing control and conditional urothelial female *Tmem63b* KO mice (Fig. 3A). A void spot analysis was also performed in conditional urothelial male *Tmem63b* KO mice and controls. Again, there was no significant difference in measures of PVS number, average PVS, total PVS, SVS number, total SVS, or the tPVS/(tPVS+tSVS) ratio in light or dark phases.

**Fig. 3.**
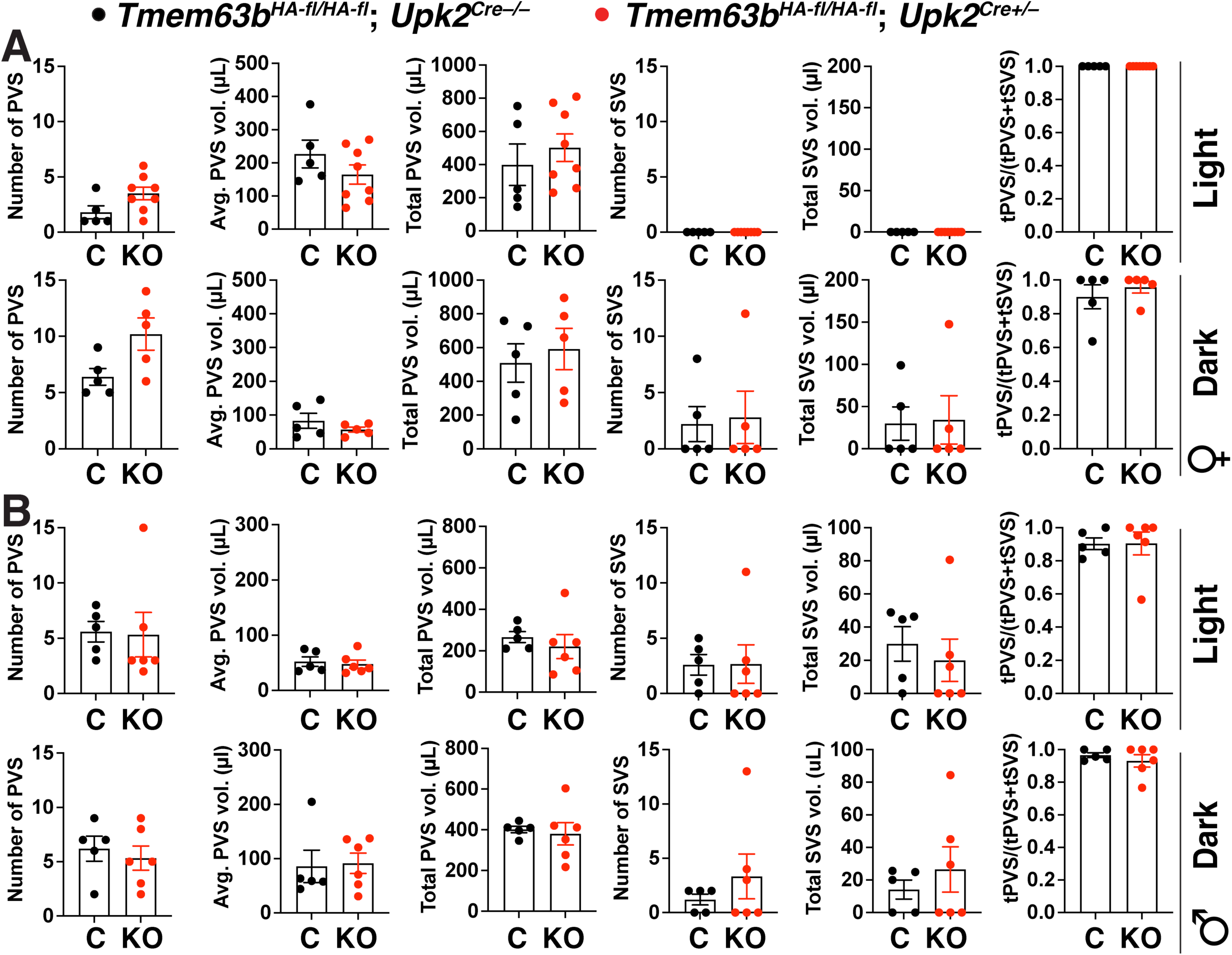
Voiding behavior in control and conditional urothelial *Tmem63b* KO male and female mice during their light and dark phases. **(A-B)** Video-monitored void spot analysis was performed in female and male *Tmem63b^HA-fl/HA-fl^*;*Upk2^Cre-/-^*(control; C) and conditional urothelial *Tmem63b^HA-fl/HA-fl^*;*Upk2^Cre+/-^*(knockout; KO) mice during the indicated light/dark phase. Measured parameters included: number of primary void spots (PVS), average number of PVS, and total volume of PVS; number of small void spots (SVS) and total volume of SVS; ratio of total PVS volume (tPVS) to total urinary output (tPVS+tSVS). Data, mean±SEM (n≥5), were analyzed using Mann-Whitney tests. No significant differences were detected.

We next assessed whether a function for *Tmem63b* would be revealed if the urothelium was stressed. In this case, we performed void spot analysis in animals treated with cyclophosphamide (CYP), an anti-cancer and anti-inflammatory drug that is rapidly converted into the toxic metabolite acrolein and excreted into urine [63–65]. Acute CYP treatment results in significant edema and swelling of the bladder lamina propria, disruption of the urothelium, and a hemorrhagic cystitis within hours of injection [66]. Relative to animal treated with vehicle (i.e., saline) alone, CYP-treated animals exhibited all of the hallmarks of an overactive bladder phenotype including a significant increase in the number of SVS (Fig. 4). The contribution of SVS volume to total urinary output increased to such an extent that the tPVS/(tPVS+tSVS) ratio fell by > 50%. These changes were accompanied by a significant decrease in number of PVS, average PVS volume, and total PVS volume. Despite these changes, there was no significant difference between the voiding behavior of control mice versus conditional urothelial *Tmem63b* KO mice in response to saline or CYP treatment (Fig. 4). On balance, we found no evidence that conditional urothelial *Tmem63b* KO mice exhibited altered voiding behavior under the conditions we tested.

**Fig. 4.**
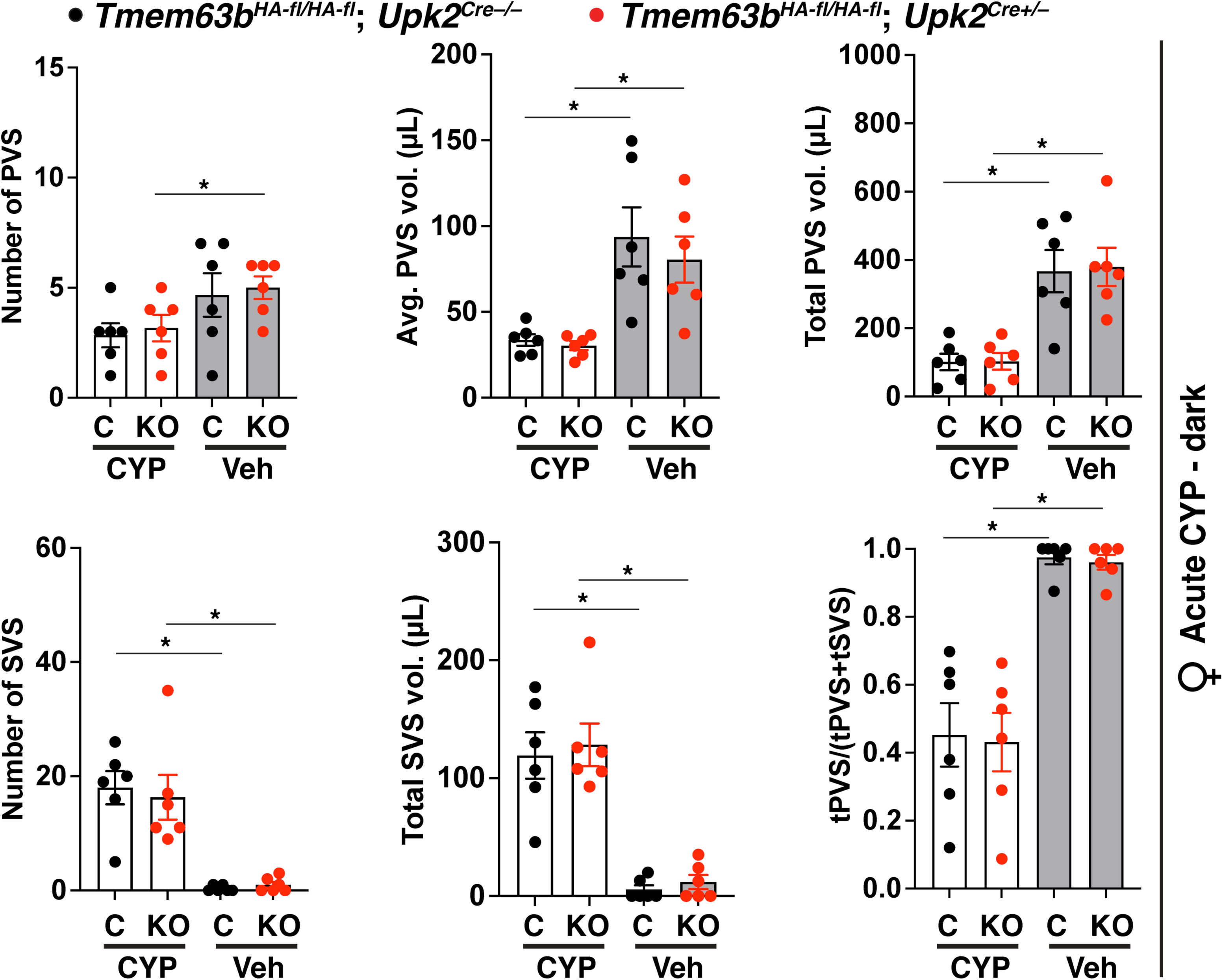
Effects of acute cyclophosphamide treatment on voiding behavior in conditional urothelial *Tmem63b* KO mice. Video-monitored void spot analysis was performed in female control *Tmem63b^HA-fl/HA-fl^*;*Upk2^Cre-/-^* (C) and conditional urothelial *Tmem63b^HA-fl/HA-fl^*;*Upk2^Cre+/-^* (KO) mice during their dark phase. Animals were monitored after injection with vehicle (Veh; saline) and subsequently after injection with 150 mg/kg of cyclophosphamide (CYP). Data are mean±SEM (n=6). Data were analyzed using two-way ANOVA with Šidák’s multiple comparisons test. Significant differences (p < 0.05) are indicated with asterisks.

### HA-TMEM63B is expressed in dorsal root ganglia neurons but loss of Tmem63b expression in DRGs does not impact voiding behavior

Additional sites of mechanotransduction in the bladder wall are the afferent processes that innervate the bladder wall and convey information about the mechanochemical status of bladder tissues to spinal cord-associated DRG neurons [3, 67–75]. FISH revealed co-expression of *Tmem63a* and *Tmem63b* in L6-S2 DRG (i.e., which include the sensory neurons that innervate the bladder)(Fig. 5A) and immunofluorescence confirmed that a subset of these DRGs expressed HA-TMEM63B (Fig. 5B). No HA signal was detected in the absence of primary antibody (Fig. 5C). Retrograde tracing studies, employing cholera toxin beta subunit (Ctx) that was injected into the bladder wall, confirmed that HA-TMEM63B was expressed in the sensory neurons that innervated the urinary bladder (Fig. 5B). As expected, none of the DRGs positioned at L4-L5 were labeled with Ctx (Fig. 5B). However, these latter neurons expressed HA-TMEM63B, indicating that TMEM63B-positive DRG neurons likely innervate other regions of the body (in the case of L4-L5, this includes the lower back, hindlimbs, and some pelvic organs). To target *Tmem63b* expression in afferent neurons, including those innervating the bladder, we used *Avil^Cre^* mice [48]. We confirmed that conditional sensory neuron *Tmem63b* KO mice (*Tmem63b^HA-fl/HA-fl^*;*Avil^Cre+/-^*) exhibited a large decrease in expression of HA-TMEM63B relative to control mice (*Tmem63b^HA-fl/HA-fl^*;*Avil^Cre-/-^*)(Fig. 5C).

**Fig. 5.**
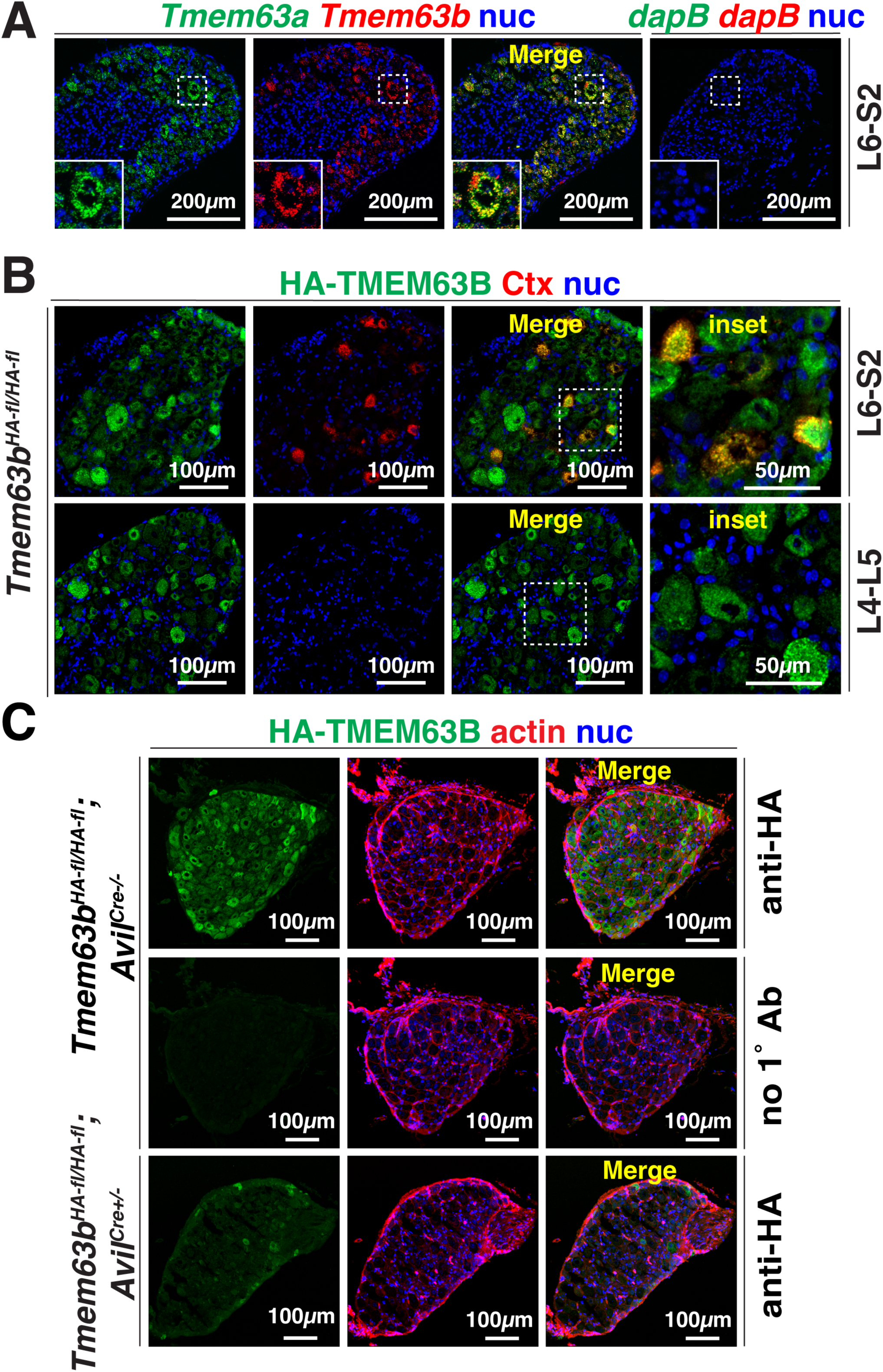
Expression of *Tmem63a*, *Tmem63b,* and HA-TMEM63B in sensory neurons. **(A)** Left-most panels: expression of *Tmem63a* and *Tmem63b* in L6-S2 DRG neurons. Right-most panel: DRG labeled with the *dapB* triple-negative control probe. The boxed regions are magnified in the insets. **(B)** HA-TMEM63B is expressed in cholera toxin (Ctx)-labeled bladder L6-S2 DRG neurons. HA-TMEM63B is also expressed in Ctx-negative L4-L5 DRG neurons. The boxed regions in the panels labeled “merge” are magnified in the right-most panels labeled “inset.” **(C)** HA-TMEM63B expression in control mice (*Tmem63b^HA-fl/HA-fl^*;*Avil^Cre-/-^*) *a*nd conditional sensory neuron KO mice *(Tmem63b^HA-fl/HA-fl^*;*Avil^Cre+/-^*). A no primary antibody control is included in the middle panel of this figure.

Both female and male control and conditional sensory neuron *Tmem63b* KO mice were subjected to void-spot analysis. Regardless of sex, or dark/light phase, we did not detect significant differences between control and conditional sensory neuron KO mice (Fig. 6A-B). CYP is reported to result in sensory neuron hypersensitivity [52, 76, 77]. Thus, we also assessed voiding behavior in CYP-treated control and conditional sensory neuron KO mice. Again, we were able to detect significant effects of CYP treatment on voiding parameters in CYP-treated versus vehicle-treated mice, but there was no significant difference in any of the measured parameters when comparing control and conditional sensory neuron KO mice (Fig. 7).

**Fig. 6.**
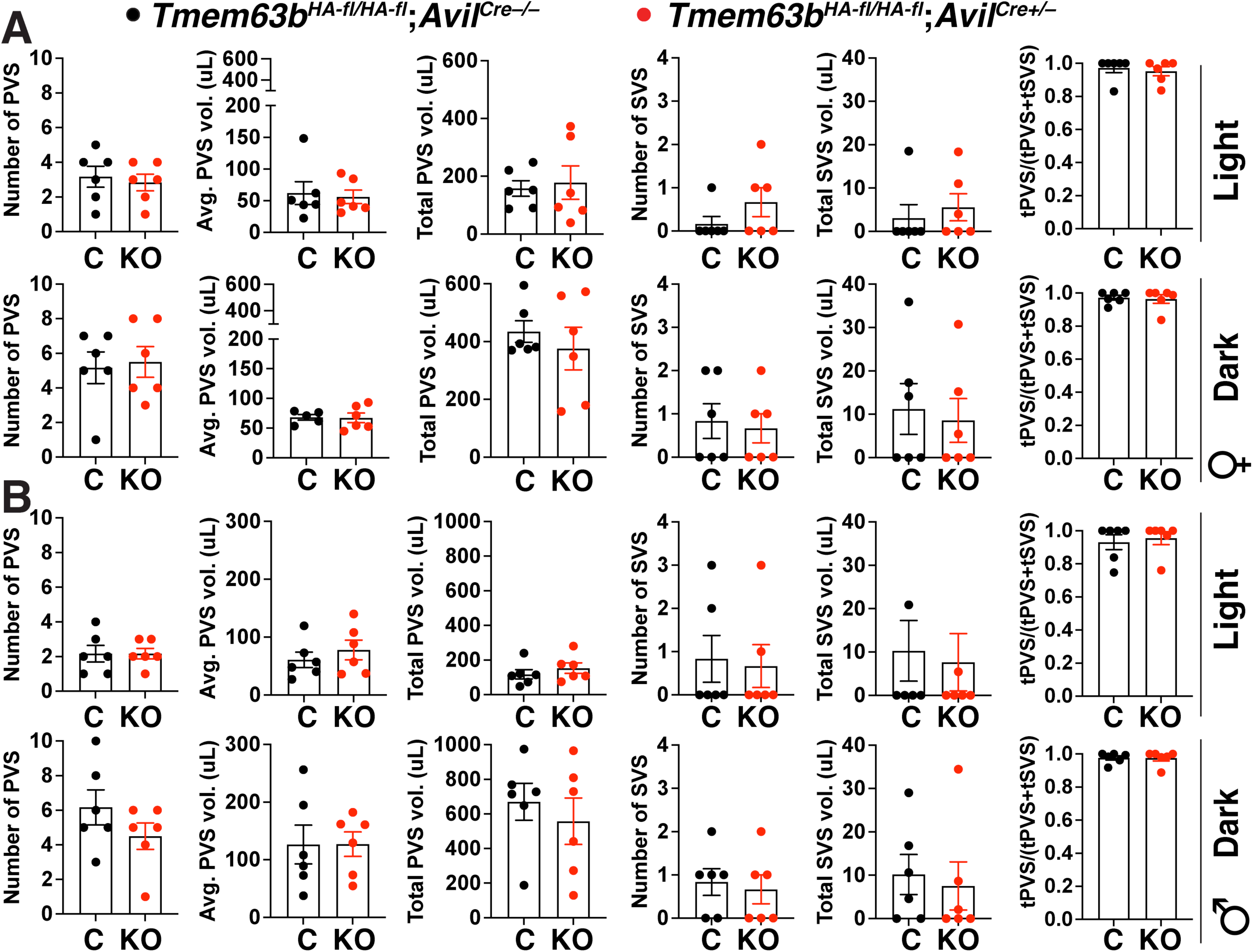
Voiding behavior in control and conditional sensory neuron *Tmem63b* KO male and female mice during their light and dark phases. **(A-B)** Video-monitored void spot analysis was performed in female and male control *Tmem63b^HA-fl/HA-fl^*;*Avil^Cre-/-^* (C) and conditional sensory neuron *Tmem63b^HA-fl/HA-fl^*;*Avil^Cre+/-^* (KO) mice during the indicated light/dark phase. Data are mean±SEM (n=6). Data were analyzed using Mann-Whitney tests. No significant differences were detected.

**Fig. 7.**
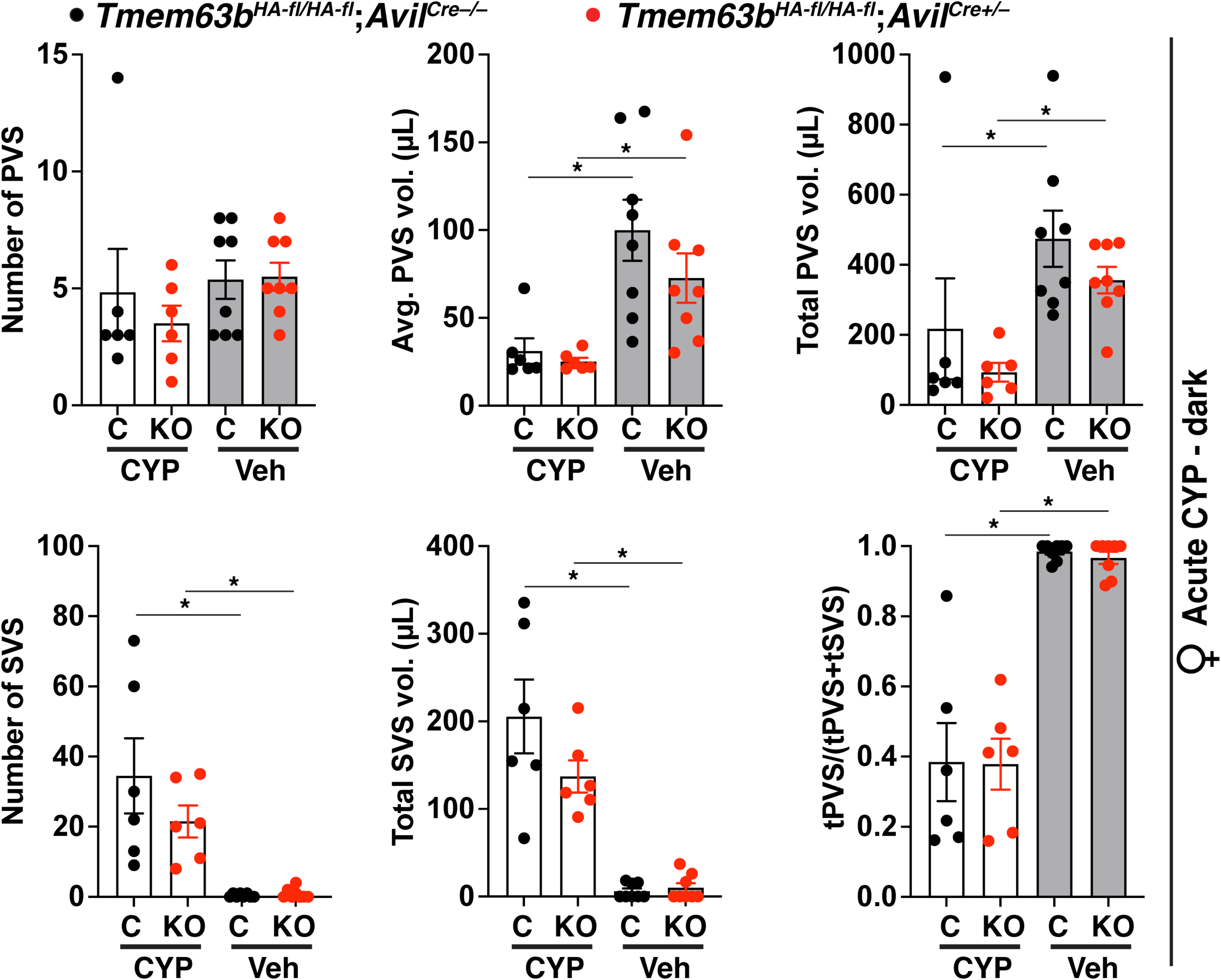
Effects of acute cyclophosphamide treatment on voiding behavior in conditional sensory neuron *Tmem63b* KO mice. Video-monitored void spot analysis was performed in female control *Tmem63b^HA-fl/HA-fl^*;*Avil^Cre-/-^* (C) and conditional sensory neuron *Tmem63b^HA-fl/HA-fl^*;*Avil^Cre+/-^*(KO) mice during their dark phase. Animals were injected with saline vehicle (Veh) or they were injected with 150 mg/kg of cyclophosphamide (CYP) prior to analysis. Data are mean±SEM (n ≥ 6). Data were analyzed using two-way ANOVA with Tukey’s multiple comparisons test. Significant differences (p < 0.05) are indicated with asterisks.

## DISCUSSION

Despite numerous reports that the urothelium and associated sensory neurons are mechanosensitive [1, 13, 78], the relevant mechanosensors and downstream mechanotransduction pathways involved in these pathways are only slowly coming into focus. Earlier evidence implicated *Itgb1* (integrin beta 1), *Trpv4*, and *Slc17a9* (vesicular nucleotide transporter) in urothelial mechanosensation [57–59]. Integrins have a well-established function in mechanotransduction [26, 79], and conditional urothelial *Itgb1* KO mice exhibit elevated mucosal ATP release (serosal release was not determined), as well as bladder overactivity and incontinence when assessed in void-spot studies [57]. The exact mechanosensory pathways affected in these mice has not yet been defined, although it is worth noting that there is increasing evidence of crosstalk between PIEZO1 signaling (see discussion below) and focal adhesions [80]. Whole animal *Trpv4* KO mice exhibit decreased mucosal ATP release (again, serosal release was not measured) and increased numbers of void spots [58]. Parsing the affected tissue types is difficult as multiple cell types in the bladder wall express *Trpv4,* as do DRGs [55, 81–83]. Furthermore, it remains controversial whether *Trpv4* functions as a mechanosensor or whether it is activated downstream of initiating mechanosensory events; however, this may depend on cell context [84, 85]. Whole animal *Slc17a9* KO mice exhibit decreased mucosal ATP release (serosal release not measured) when the bladder is filled with small fluid volumes [59]. In addition, these mice exhibit decreased bladder compliance and have an increased tendency to void away from their corners when analyzed in void-spot assays. *Slc17a9* is expressed by the urothelium as well as sensory neurons [59, 86]. Thus, the underlying target(s) of the KO phenotype is not yet clear.

More recent data implicate the well described mechanosensory proteins PIEZO1 and PIEZO2 in urothelial and afferent function [27]. In our hands, individual conditional urothelial *Piezo1* or *Piezo2* KO mice exhibit a limited bladder phenotype [2]. In contrast, double conditional urothelial *Piezo1/2* KO are characterized by an almost complete loss of urothelial serosal ATP release and altered bladder and voiding behavior that varies in a sex- and circadian rhythm-dependent manner [2]. Worth noting though, Marshall *et al.* report that conditional urothelial *Piezo2* KO mice exhibit altered bladder and sphincter function, but they did not measure the impact of *Piezo2* KO on urothelial mechanotransduction or voiding behavior [3]. One would predict that ablating urothelial pathways for mechanotransduction would lead to a hypoactive bladder. Instead, conditional urothelial *Itgb1KO mice, Trpv4* KO mice, and double *Piezo1/2* KO mice have an overactive bladder phenotype (characterized by large numbers of small voiding events) indicating that urothelial mechanotransduction may play a role in suppressing voiding function and not simply stimulating it [2]. Sensory neurons are also known to express *Piezo2* and Marshall *et al.* report that conditional sensory neuron *Piezo2* KO mice exhibit reduced mechanosensation and have a bladder phenotype that includes increased intercontraction interval, larger voiding pressures, and decreased urethral contraction activity [3]. Strikingly, *PIEZO2*-deficient patients also exhibit voiding defects including decreased voiding frequency, in some instances coupled with urge incontinence [3]. While a role for urothelial PIEZO2 expression cannot be excluded in these patients, one likely target is sensory neurons as *PIEZO2*-deficient individuals and mice also suffer from defects in proprioception [87–89].

Our current studies were focused on identifying a role for TMEM63B in voiding behavior. This follows up on recent studies that implicate TMEM63B in mechano- and osmo-sensation across a variety of organs, as well as our recent evidence that HA-TMEM63B is expressed within the upper and lower urinary tracts including the urothelium lining the renal pelvis, ureters, bladder, and upper urethra [50]. Our current studies further reveal that the urothelium expresses both *Tmem63a* and *Tmem63b*, and that DRG neurons, including those innervating the bladder, express HA-TMEM63B (as well as *Tmem63a* and *Tmem63b*). We further generated conditional urothelial *Tmem63b* KO mice as well as conditional sensory neuron *Tmem63b* KO mice. However, using a well-documented screening tool for defects in lower urinary tract function, the void-spot assay [2, 3, 57–59], we were unable to detect a phenotype resulting from loss of urothelial or sensory neuron *Tmem63b* expression. This was despite efforts to look for sex-specific differences or variations that could be ascribed to circadian effects. We also assessed whether the overactive bladder phenotype that results from treatment with CYP would be affected by loss of *Tmem63b* expression. However, we could not detect a difference.

One could draw the conclusion that *Tmem63b* plays no role in normal bladder voiding behavior and responses to acute CYP treatment; however, there are several caveats worth considering. The first, is that we may not have found the right conditions to reveal a role for *Tmem63b*. Thus, by altering urine composition (e.g., making it hypertonic or hypotonic) or by exposing the bladder to bacterial infections, or chronic disease states such as partial outlet obstruction or spinal cord injury, we may reveal a role for *Tmem63b*. The second caveat is that the phenotype resulting from loss of *Tmem63b* expression in the urothelium (or sensory neurons) may be complex and involve compensatory functional changes, which may not be revealed by void spot assays. The third caveat is that there may be compensatory changes in gene expression that accompany knockout of *Tmem63b*. We noted that the urothelium and sensory neurons express both *Tmem63a* and *Tmem63b*. Thus, only a double *Tmem63a/Tmem63b* KO mouse may reveal a phenotype. Such a possibility has been reported for *Tmem63a/b*-dependent mechano/osmosensation in lung epithelia [40], and as noted above, we previously reported that a bladder phenotype was only revealed in conditional urothelial *Piezo1/2* double KO mice and not individual conditional urothelial *Piezo1* or *Piezo2* KO animals [2]. An obvious next step will be to generate double *Tmem63a/b* KO animals and define their phenotype.

In sum, our current studies indicate that despite expression of TMEM63B in the urothelium and sensory neurons, loss of *Tmem63b* expression in these tissues does not result in an obvious phenotype when assessing voiding behavior. However, we cannot exclude the possibility that *Tmem63b* function will be revealed in other physiological/pathological states or when expression of both *Tmem63b* and *Tmem63a* are simultaneously knocked out.

## ACKNOWLEDGEMENTS

This work was supported by grants from the National institutes of Health including R01DK119183 (to GA and MDC), R01DK129473 (to GA), a pilot project grant supported by P30DK079307 (to MGD), and by the Pittsburgh Center for Kidney Research KIDNIT imaging core (U54DK137329). The work was also supported by grants from the National Natural Science Foundation of China 32330044 (to YSS). The Leica Stellaris confocal used in this study was funded in large part by S10OD028596 (to GA).

## AUTHOR CONTRIBUTIONS

Conceived and designed research: MD, WR, DC, GA, MDC, YSS

Performed experiments, analyzed data, and interpreted results of experiments: MD, WR, DC, TP, MDC, GA

Prepared figures: DC, GA

Drafted manuscript: MD, MDC, GA

Edited and revised manuscript: MD, MDC, YSS, GA

Approved final version of manuscript: MD, WR, DC, TP, MDC, YSS, GA

## DATA availability

The data that support the findings of this study are available from the corresponding author upon reasonable request.

## REFERENCES

1. Dalghi MG, Montalbetti N, Carattino MD, Apodaca G. The Urothelium: Life in a Liquid Environment. Physiol Rev. 2020;100(4):1621–705. Epub 20200319. doi: 10.1152/physrev.00041.2019. PubMed PMID: 32191559; PubMed Central PMCID: PMCPMC7717127.

2. Dalghi MG, Ruiz WG, Clayton DR, Montalbetti N, Daugherty SL, Beckel JM, et al. Functional roles for PIEZO1 and PIEZO2 in urothelial mechanotransduction and lower urinary tract interoception. Journal of Clinical Invesitgation Insight. 2021;6(19). Epub 20211008. doi: 10.1172/jci.insight.152984. PubMed PMID: 34464353; PubMed Central PMCID: PMCPMC8525643.

3. Marshall KL, Saade D, Ghitani N, Coombs AM, Szczot M, Keller J, et al. PIEZO2 in sensory neurons and urothelial cells coordinates urination. Nature. 2020;588(7837):290-5. Epub 20201014. doi: 10.1038/s41586-020-2830-7. PubMed PMID: 33057202; PubMed Central PMCID: PMCPMC7725878.

4. Dalghi MG, Clayton DR, Ruiz WG, Al-Bataineh MM, Satlin LM, Kleyman TR, et al. Expression and distribution of PIEZO1 in the mouse urinary tract. Am J Physiol Renal Physiol. 2019;317(2):F303–F21. Epub 2019/06/06. doi: 10.1152/ajprenal.00214.2019. PubMed PMID: 31166705.

5. Persson K, Sando JJ, Tuttle JB, Steers WD. Protein kinase C in cyclic stretch-induced nerve growth factor production by urinary tract smooth muscle cells. Am J Physiol. 1995;269(4 Pt 1):C1018-24. doi: 10.1152/ajpcell.1995.269.4.C1018. PubMed PMID: 7485441.

6. Wellner MC, Isenberg G. Properties of stretch-activated channels in myocytes from the guinea-pig urinary bladder. J Physiol. 1993;466:213–27. Epub 1993/07/01. PubMed PMID: 7692040; PubMed Central PMCID: PMCPMC1175475.

7. Pineda RH, Hypolite J, Lee S, Carrasco A, Jr., Iguchi N, Meacham RB, et al. Altered detrusor contractility and voiding patterns in mice lacking the mechanosensitive TREK-1 channel. BMC Urol. 2019;19(1):40. Epub 20190521. doi: 10.1186/s12894-019-0475-3. PubMed PMID: 31113422; PubMed Central PMCID: PMCPMC6528348.

8. Roberts MWG, Sui G, Wu R, Rong W, Wildman S, Montgomery B, et al. TRPV4 receptor as a functional sensory molecule in bladder urothelium: Stretch-independent, tissue-specific actions and pathological implications. FASEB J. 2020;34(1):263–86. Epub 2020/01/10. doi: 10.1096/fj.201900961RR. PubMed PMID: 31914645.

9. Liu Q, Sun B, Zhao J, Wang Q, An F, Hu X, et al. Increased Piezo1 channel activity in interstitial Cajal-like cells induces bladder hyperactivity by functionally interacting with NCX1 in rats with cyclophosphamide-induced cystitis. Exp Mol Med. 2018;50(5):1–16. Epub 20180507. doi: 10.1038/s12276-018-0088-z. PubMed PMID: 29735991; PubMed Central PMCID: PMCPMC5938236.

10. Apodaca G, Balestreire E, Birder LA. The uroepithelial-associated sensory web. Kidney International. 2007;72:1057–64.

11. Birder L, Andersson KE. Urothelial signaling. Physiological Reviews. 2013;93(2):653-80. doi: 10.1152/physrev.00030.2012. PubMed PMID: 23589830; PubMed Central PMCID: PMCPMC3768101.

12. Birder LA. Urinary bladder, cystitis and nerve/urothelial interactions. Autonomic Neuroscience. 2014;182:89–94. Epub 2014/01/15. doi: 10.1016/j.autneu.2013.12.005. PubMed PMID: 24412640; PubMed Central PMCID: PMCPMC3989437.

13. Li X, Hu J, Yin P, Liu L, Chen Y. Mechanotransduction in the urothelium: ATP signalling and mechanoreceptors. Heliyon. 2023;9(9):e19427. Epub 20230823. doi: 10.1016/j.heliyon.2023.e19427. PubMed PMID: 37674847; PubMed Central PMCID: PMCPMC10477517.

14. Goodman MB, Haswell ES, Vasquez V. Mechanosensitive membrane proteins: Usual and unusual suspects in mediating mechanotransduction. The Journal of general physiology. 2023;155(3). Epub 20230125. doi: 10.1085/jgp.202213248. PubMed PMID: 36696153; PubMed Central PMCID: PMCPMC9930137.

15. Ingber DE. Cellular mechanotransduction: putting all the pieces together again. Faseb J. 2006;20(7):811–27. PubMed PMID: 16675838.

16. Martino F, Perestrelo AR, Vinarsky V, Pagliari S, Forte G. Cellular Mechanotransduction: From Tension to Function. Front Physiol. 2018;9:824. Epub 20180705. doi: 10.3389/fphys.2018.00824. PubMed PMID: 30026699; PubMed Central PMCID: PMCPMC6041413.

17. Hayakawa K, Tatsumi H, Sokabe M. Actin filaments function as a tension sensor by tension-dependent binding of cofilin to the filament. J Cell Biol. 2011;195(5):721–7. doi: 10.1083/jcb.201102039. PubMed PMID: 22123860; PubMed Central PMCID: PMCPMC3257564.

18. van Bodegraven EJ, Etienne-Manneville S. Intermediate Filaments from Tissue Integrity to Single Molecule Mechanics. Cells. 2021;10(8). Epub 20210727. doi: 10.3390/cells10081905. PubMed PMID: 34440673; PubMed Central PMCID: PMCPMC8392029.

19. Hatzfeld M, Keil R, Magin TM. Desmosomes and Intermediate Filaments: Their Consequences for Tissue Mechanics. Cold Spring Harbor perspectives in biology. 2017;9(6). Epub 20170601. doi: 10.1101/cshperspect.a029157. PubMed PMID: 28096266; PubMed Central PMCID: PMCPMC5453391.

20. Seetharaman S, Vianay B, Roca V, Farrugia AJ, De Pascalis C, Boeda B, et al. Microtubules tune mechanosensitive cell responses. Nat Mater. 2022;21(3):366–77. Epub 20211018. doi: 10.1038/s41563-021-01108-x. PubMed PMID: 34663953.

21. Hoffman BD, Yap AS. Towards a Dynamic Understanding of Cadherin-Based Mechanobiology. Trends Cell Biol. 2015;25(12):803–14. Epub 2015/11/02. doi: 10.1016/j.tcb.2015.09.008. PubMed PMID: 26519989.

22. Bachmann M, Kukkurainen S, Hytönen VP, Wehrle-Haller B. Cell Adhesion by Integrins. Physiol Rev. 2019;99(4):1655–99. doi: 10.1152/physrev.00036.2018. PubMed PMID: 31313981.

23. Charras G, Yap AS. Tensile Forces and Mechanotransduction at Cell-Cell Junctions. Curr Biol. 2018;28(8):R445–R57. Epub 2018/04/25. doi: 10.1016/j.cub.2018.02.003. PubMed PMID: 29689229.

24. Angulo-Urarte A, van der Wal T, Huveneers S. Cell-cell junctions as sensors and transducers of mechanical forces. Biochim Biophys Acta Biomembr. 2020;1862(9):183316. Epub 20200428. doi: 10.1016/j.bbamem.2020.183316. PubMed PMID: 32360073.

25. Wilde C, Mitgau J, Suchy T, Schoneberg T, Liebscher I. Translating the force-mechano-sensing GPCRs. Am J Physiol Cell Physiol. 2022;322(6):C1047–C60. Epub 20220413. doi: 10.1152/ajpcell.00465.2021. PubMed PMID: 35417266.

26. Kolasangiani R, Bidone TC, Schwartz MA. Integrin Conformational Dynamics and Mechanotransduction. Cells. 2022;11(22). Epub 20221112. doi: 10.3390/cells11223584. PubMed PMID: 36429013; PubMed Central PMCID: PMCPMC9688440.

27. Kefauver JM, Ward AB, Patapoutian A. Discoveries in structure and physiology of mechanically activated ion channels. Nature. 2020;587(7835):567-76. Epub 2020/11/27. doi: 10.1038/s41586-020-2933-1. PubMed PMID: 33239794.

28. Zhao X, Yan X, Liu Y, Zhang P, Ni X. Co-expression of mouse TMEM63A, TMEM63B and TMEM63C confers hyperosmolarity activated ion currents in HEK293 cells. Cell Biochem Funct. 2016;34(4):238-41. Epub 20160404. doi: 10.1002/cbf.3185. PubMed PMID: 27045885.

29. Du H, Ye C, Wu D, Zang YY, Zhang L, Chen C, et al. The Cation Channel TMEM63B Is an Osmosensor Required for Hearing. Cell Rep. 2020;31(5):107596. doi: 10.1016/j.celrep.2020.107596. PubMed PMID: 32375046.

30. Murthy SE, Dubin AE, Whitwam T, Jojoa-Cruz S, Cahalan SM, Mousavi SAR, et al. OSCA/TMEM63 are an Evolutionarily Conserved Family of Mechanically Activated Ion Channels. Elife. 2018;7. Epub 20181101. doi: 10.7554/eLife.41844. PubMed PMID: 30382938; PubMed Central PMCID: PMCPMC6235560.

31. Zheng W, Rawson S, Shen Z, Tamilselvan E, Smith HE, Halford J, et al. TMEM63 proteins function as monomeric high-threshold mechanosensitive ion channels. Neuron. 2023;111(20):3195–210 e7. Epub 20230804. doi: 10.1016/j.neuron.2023.07.006. PubMed PMID: 37543036; PubMed Central PMCID: PMCPMC10592209.

32. Li S, Li B, Gao L, Wang J, Yan Z. Humidity response in Drosophila olfactory sensory neurons requires the mechanosensitive channel TMEM63. Nature communications. 2022;13(1):3814. Epub 20220702. doi: 10.1038/s41467-022-31253-z. PubMed PMID: 35780140; PubMed Central PMCID: PMCPMC9250499.

33. Li Q, Montell C. Mechanism for food texture preference based on grittiness. Current biology : CB. 2021;31(9):1850–61 e6. Epub 20210302. doi: 10.1016/j.cub.2021.02.007. PubMed PMID: 33657409; PubMed Central PMCID: PMCPMC8119346.

34. Li K, Guo Y, Wang Y, Zhu R, Chen W, Cheng T, et al. Drosophila TMEM63 and mouse TMEM63A are lysosomal mechanosensory ion channels. Nature cell biology. 2024;26(3):393–403. Epub 20240222. doi: 10.1038/s41556-024-01353-7. PubMed PMID: 38388853; PubMed Central PMCID: PMCPMC10940159.

35. Pu S, Wu Y, Tong F, Du WJ, Liu S, Yang H, et al. Mechanosensitive Ion Channel TMEM63A Gangs Up with Local Macrophages to Modulate Chronic Post-amputation Pain. Neurosci Bull. 2023;39(2):177–93. Epub 20220712. doi: 10.1007/s12264-022-00910-0. PubMed PMID: 35821338; PubMed Central PMCID: PMCPMC9905372.

36. Yan H, Helman G, Murthy SE, Ji H, Crawford J, Kubisiak T, et al. Heterozygous Variants in the Mechanosensitive Ion Channel TMEM63A Result in Transient Hypomyelination during Infancy. American journal of human genetics. 2019;105(5):996–1004. Epub 20191003. doi: 10.1016/j.ajhg.2019.09.011. PubMed PMID: 31587869; PubMed Central PMCID: PMCPMC6848986.

37. Yan H, Ji H, Kubisiak T, Wu Y, Xiao J, Gu Q, et al. Genetic analysis of 20 patients with hypomyelinating leukodystrophy by trio-based whole-exome sequencing. J Hum Genet. 2021;66(8):761–8. Epub 20210218. doi: 10.1038/s10038-020-00896-5. PubMed PMID: 33597727; PubMed Central PMCID: PMCPMC8310791.

38. Fukumura S, Hiraide T, Yamamoto A, Tsuchida K, Aoto K, Nakashima M, et al. A novel de novo TMEM63A variant in a patient with severe hypomyelination and global developmental delay. Brain Dev. 2022;44(2):178–83. Epub 20210928. doi: 10.1016/j.braindev.2021.09.006. PubMed PMID: 34598833.

39. Wang YY, Wu D, Zhan Y, Li F, Zang YY, Teng XY, et al. Cation Channel TMEM63A Autonomously Facilitates Oligodendrocyte Differentiation at an Early Stage. Neurosci Bull. 2025. Epub 20250221. doi: 10.1007/s12264-024-01338-4. PubMed PMID: 39982638.

40. Chen GL, Li JY, Chen X, Liu JW, Zhang Q, Liu JY, et al. Mechanosensitive channels TMEM63A and TMEM63B mediate lung inflation-induced surfactant secretion. J Clin Invest. 2024;134(5). Epub 20241221. doi: 10.1172/JCI174508. PubMed PMID: 38127458; PubMed Central PMCID: PMCPMC10904053.

41. Yang G, Jia M, Li G, Zang YY, Chen YY, Wang YY, et al. TMEM63B channel is the osmosensor required for thirst drive of interoceptive neurons. Cell Discov. 2024;10(1):1. Epub 20240103. doi: 10.1038/s41421-023-00628-x. PubMed PMID: 38172113; PubMed Central PMCID: PMCPMC10764952.

42. Tu JJ, Ye C, Teng XY, Zang YY, Sun XY, Chen S, et al. Osmosensor TMEM63B facilitates insulin secretion in pancreatic beta-cells. Sci China Life Sci. 2025. Epub 20250220. doi: 10.1007/s11427-024-2833-3. PubMed PMID: 39985646.

43. Ye C, Zhang TZ, Zang YY, Shi YS, Wan G. TMEM63B regulates postnatal development of cochlear sensory epithelia via thyroid hormone signaling. J Genet Genomics. 2024;51(6):673–6. Epub 20231227. doi: 10.1016/j.jgg.2023.12.006. PubMed PMID: 38157934.

44. Vetro A, Pelorosso C, Balestrini S, Masi A, Hambleton S, Argilli E, et al. Stretch-activated ion channel TMEM63B associates with developmental and epileptic encephalopathies and progressive neurodegeneration. American journal of human genetics. 2023;110(8):1356–76. Epub 20230707. doi: 10.1016/j.ajhg.2023.06.008. PubMed PMID: 37421948; PubMed Central PMCID: PMCPMC10432263.

45. Tabara LC, Al-Salmi F, Maroofian R, Al-Futaisi AM, Al-Murshedi F, Kennedy J, et al. TMEM63C mutations cause mitochondrial morphology defects and underlie hereditary spastic paraplegia. Brain. 2022;145(9):3095–107. doi: 10.1093/brain/awac123. PubMed PMID: 35718349; PubMed Central PMCID: PMCPMC9473353.

46. Schulz A, Muller NV, van de Lest NA, Eisenreich A, Schmidbauer M, Barysenka A, et al. Analysis of the genomic architecture of a complex trait locus in hypertensive rat models links Tmem63c to kidney damage. Elife. 2019;8. Epub 20190322. doi: 10.7554/eLife.42068. PubMed PMID: 30900988; PubMed Central PMCID: PMCPMC6478434.

47. Ayala de la Pena F, Kanasaki K, Kanasaki M, Tangirala N, Maeda G, Kalluri R. Loss of p53 and acquisition of angiogenic microRNA profile are insufficient to facilitate progression of bladder urothelial carcinoma in situ to invasive carcinoma. J Biol Chem. 2011;286(23):20778–87. Epub 20110309. doi: 10.1074/jbc.M110.198069. PubMed PMID: 21388952; PubMed Central PMCID: PMCPMC3121487.

48. Zhou X, Wang L, Hasegawa H, Amin P, Han BX, Kaneko S, et al. Deletion of PIK3C3/Vps34 in sensory neurons causes rapid neurodegeneration by disrupting the endosomal but not the autophagic pathway. Proc Natl Acad Sci U S A. 2010;107(20):9424–9. Epub 20100503. doi: 10.1073/pnas.0914725107. PubMed PMID: 20439739; PubMed Central PMCID: PMCPMC2889054.

49. Truett GE, Heeger P, Mynatt RL, Truett AA, Walker JA, Warman ML. Preparation of PCR-quality mouse genomic DNA with hot sodium hydroxide and tris (HotSHOT). Biotechniques. 2000;29(1):52, 4. doi: 10.2144/00291bm09. PubMed PMID: 10907076.

50. Dalghi MG, DuRie E, Ruiz WG, Clayton DR, Montalbetti N, Mutchler SB, et al. Expression and localization of the mechanosensitive/osmosensitive ion channel TMEM63B in the mouse urinary tract. Physiol Rep. 2024;12(9):e16043. doi: 10.14814/phy2.16043. PubMed PMID: 38724885; PubMed Central PMCID: PMCPMC11082094.

51. Montalbetti N, Rooney JG, Marciszyn AL, Carattino MD. ASIC3 fine-tunes bladder sensory signaling. Am J Physiol Renal Physiol. 2018;315(4):F870-f9. Epub 20180321. doi: 10.1152/ajprenal.00630.2017. PubMed PMID: 29561183; PubMed Central PMCID: PMCPMC6230751.

52. Montalbetti N, Carattino MD. Acid-sensing ion channels modulate bladder nociception. Am J Physiol Renal Physiol. 2021;321(5):F587-f99. Epub 20210913. doi: 10.1152/ajprenal.00302.2021. PubMed PMID: 34514879; PubMed Central PMCID: PMCPMC8813206.

53. Dalghi MG, Montalbetti N, Wheeler TB, Apodaca G, Carattino MD. Real-Time Void Spot Assay. J Vis Exp. 2023;(192). Epub 20230210. doi: 10.3791/64621. PubMed PMID: 36847378.

54. Hill WG, Zeidel ML, Bjorling DE, Vezina CM. Void spot assay: recommendations on the use of a simple micturition assay for mice. Am J Physiol Renal Physiol. 2018;315(5):F1422–F9. Epub 20180829. doi: 10.1152/ajprenal.00350.2018. PubMed PMID: 30156116; PubMed Central PMCID: PMCPMC6293303.

55. Yu Z, Liao J, Chen Y, Zou C, Zhang H, Cheng J, et al. Single-Cell Transcriptomic Map of the Human and Mouse Bladders. J Am Soc Nephrol. 2019;30(11):2159–76. Epub 2019/08/30. doi: 10.1681/ASN.2019040335. PubMed PMID: 31462402; PubMed Central PMCID: PMCPMC6830796.

56. Mo L, Cheng J, Lee EY, Sun TT, Wu XR. Gene deletion in urothelium by specific expression of Cre recombinase. Am J Physiol Renal Physiol. 2005;289(3):F562–8. Epub 2005/04/21. doi: 10.1152/ajprenal.00368.2004. PubMed PMID: 15840768.

57. Kanasaki K, Yu W, von Bodungen M, Larigakis JD, Kanasaki M, Ayala de la Pena F, et al. Loss of beta1-integrin from urothelium results in overactive bladder and incontinence in mice: a mechanosensory rather than structural phenotype. FASEB J. 2013;27(5):1950–61. doi: 10.1096/fj.12-223404. PubMed PMID: 23395910; PubMed Central PMCID: PMCPMC3633821.

58. Gevaert T, Vriens J, Segal A, Everaerts W, Roskams T, Talavera K, et al. Deletion of the transient receptor potential cation channel TRPV4 impairs murine bladder voiding. J Clin Invest. 2007;117(11):3453–62. PubMed PMID: 17948126.

59. Nakagomi H, Yoshiyama M, Mochizuki T, Miyamoto T, Komatsu R, Imura Y, et al. Urothelial ATP exocytosis: regulation of bladder compliance in the urine storage phase. Sci Rep. 2016;6:29761. doi: 10.1038/srep29761. PubMed PMID: 27412485; PubMed Central PMCID: PMCPMC4944198.

60. Montalbetti N, Dalghi MG, Bastacky SI, Clayton DR, Ruiz WG, Apodaca G, et al. Bladder infection with uropathogenic Escherichia coli increases the excitability of afferent neurons. Am J Physiol Renal Physiol. 2022;322(1):F1-F13. Epub 20211115. doi: 10.1152/ajprenal.00167.2021. PubMed PMID: 34779263; PubMed Central PMCID: PMCPMC8698541.

61. Wood R, Eichel L, Messing EM, Schwarz E. Automated noninvasive measurement of cyclophosphamide-induced changes in murine voiding frequency and volume. J Urol. 2001;165(2):653–9. doi: 10.1097/00005392-200102000-00089. PubMed PMID: 11176453.

62. Rajandram R, Ong TA, Razack AH, MacIver B, Zeidel M, Yu W. Intact urothelial barrier function in a mouse model of ketamine-induced voiding dysfunction. Am J Physiol Renal Physiol. 2016;310(9):F885-94. Epub 2016/02/26. doi: 10.1152/ajprenal.00483.2015. PubMed PMID: 26911853; PubMed Central PMCID: PMCPMC4867311.

63. Cox PJ. Cyclophosphamide cystitis--identification of acrolein as the causative agent. Biochem Pharmacol. 1979;28(13):2045–9. PubMed PMID: 475846.

64. Ahlmann M, Hempel G. The effect of cyclophosphamide on the immune system: implications for clinical cancer therapy. Cancer Chemother Pharmacol. 2016;78(4):661–71. Epub 2016/09/21. doi: 10.1007/s00280-016-3152-1. PubMed PMID: 27646791.

65. Brock N, Stekar J, Pohl J, Niemeyer U, Scheffler G. Acrolein, the causative factor of urotoxic side-effects of cyclophosphamide, ifosfamide, trofosfamide and sufosfamide. Arzneimittelforschung. 1979;29(4):659–61. PubMed PMID: 114192.

66. Haldar S, Dru C, Bhowmick NA. Mechanisms of hemorrhagic cystitis. Am J Clin Exp Urol. 2014;2(3):199–208. Epub 20141002. PubMed PMID: 25374922; PubMed Central PMCID: PMCPMC4219308.

67. Floyd K, Hick VE, Morrison JF. Mechanosensitive afferent units in the hypogastric nerve of the cat. J Physiol. 1976;259(2):457–71. Epub 1976/07/01. doi: 10.1113/jphysiol.1976.sp011476. PubMed PMID: 986462; PubMed Central PMCID: PMCPMC1309039.

68. Bahns E, Ernsberger U, Janig W, Nelke A. Functional characteristics of lumbar visceral afferent fibres from the urinary bladder and the urethra in the cat. Pflugers Arch. 1986;407(5):510–8. Epub 1986/11/01. doi: 10.1007/bf00657509. PubMed PMID: 3786110.

69. Sengupta JN, Gebhart GF. Mechanosensitive properties of pelvic nerve afferent fibers innervating the urinary bladder of the rat. J Neurophysiol. 1994;72(5):2420–30. Epub 1994/11/01. doi: 10.1152/jn.1994.72.5.2420. PubMed PMID: 7884468.

70. Wen J, Morrison JF. The effects of high urinary potassium concentration on pelvic nerve mechanoreceptors and ’silent’ afferents from the rat bladder. Adv Exp Med Biol. 1995;385:237–9. Epub 1995/01/01. doi: 10.1007/978-1-4899-1585-6_29. PubMed PMID: 8571836.

71. Dmitrieva N, McMahon SB. Sensitisation of visceral afferents by nerve growth factor in the adult rat. Pain. 1996;66(1):87–97. Epub 1996/07/01. doi: 10.1016/0304-3959(96)02993-4. PubMed PMID: 8857635.

72. Kanai A, Andersson KE. Bladder afferent signaling: recent findings. J Urol. 2010;183(4):1288–95. doi: 10.1016/j.juro.2009.12.060. PubMed PMID: 20171668; PubMed Central PMCID: PMCPMC3686308.

73. Umans BD, Liberles SD. Neural Sensing of Organ Volume. Trends in neurosciences. 2018;41(12):911–24. Epub 2018/08/26. doi: 10.1016/j.tins.2018.07.008. PubMed PMID: 30143276; PubMed Central PMCID: PMCPMC6252275.

74. Gabella G. Afferent nerve fibres in the wall of the rat urinary bladder. Cell Tissue Res. 2019;376(1):25–35. Epub 20181207. doi: 10.1007/s00441-018-2965-0. PubMed PMID: 30523406.

75. Heppner TJ, Hennig GW, Nelson MT, Herrera GM. Afferent nerve activity in a mouse model increases with faster bladder filling rates in vitro, but voiding behavior remains unaltered in vivo. Am J Physiol Regul Integr Comp Physiol. 2022;323(5):R682-R93. Epub 20220919. doi: 10.1152/ajpregu.00156.2022. PubMed PMID: 36121145; PubMed Central PMCID: PMCPMC9602904.

76. Dang K, Lamb K, Cohen M, Bielefeldt K, Gebhart GF. Cyclophosphamide-induced bladder inflammation sensitizes and enhances P2X receptor function in rat bladder sensory neurons. J Neurophysiol. 2008;99(1):49–59. Epub 2007/10/26. doi: 10.1152/jn.00211.2007. PubMed PMID: 17959738; PubMed Central PMCID: PMCPMC2659400.

77. Yoshimura N, de Groat WC. Increased excitability of afferent neurons innervating rat urinary bladder after chronic bladder inflammation. J Neurosci. 1999;19(11):4644–53. PubMed PMID: 10341262.

78. Janssen DAW, Schalken JA, Heesakkers J. Urothelium update: how the bladder mucosa measures bladder filling. Acta Physiol (Oxf). 2017;220(2):201–17. doi: 10.1111/apha.12824. PubMed PMID: 27804256.

79. Katoh K. Integrin and Its Associated Proteins as a Mediator for Mechano-Signal Transduction. Biomolecules. 2025;15(2). Epub 20250123. doi: 10.3390/biom15020166. PubMed PMID: 40001469; PubMed Central PMCID: PMCPMC11853369.

80. Cheng D, Wang J, Yao M, Cox CD. Joining forces: crosstalk between mechanosensitive PIEZO1 ion channels and integrin-mediated focal adhesions. Biochem Soc Trans. 2023;51(5):1897–906. doi: 10.1042/BST20230042. PubMed PMID: 37772664.

81. Sidwell AB, McClintock C, Beca KI, Campbell SE, Girard BM, Vizzard MA. Repeated variate stress increased voiding frequency and altered TrpV1 and TrpV4 transcript expression in lower urinary tract (LUT) pathways in female mice. Front Urol. 2023;2. Epub 20230125. doi: 10.3389/fruro.2022.1086179. PubMed PMID: 37692906; PubMed Central PMCID: PMCPMC10492642.

82. Girard BM, Merrill L, Malley S, Vizzard MA. Increased TRPV4 expression in urinary bladder and lumbosacral dorsal root ganglia in mice with chronic overexpression of NGF in urothelium. J Mol Neurosci. 2013;51(2):602–14. Epub 20130521. doi: 10.1007/s12031-013-0033-5. PubMed PMID: 23690258; PubMed Central PMCID: PMCPMC3779511.

83. Thorneloe KS, Sulpizio AC, Lin Z, Figueroa DJ, Clouse AK, McCafferty GP, et al. N- ((1S)-1-{[4-((2S)-2-{[(2,4-dichlorophenyl)sulfonyl]amino}-3-hydroxypropanoyl)-1 -piperazinyl]carbonyl}-3-methylbutyl)-1-benzothiophene-2-carboxamide (GSK1016790A), a novel and potent transient receptor potential vanilloid 4 channel agonist induces urinary bladder contraction and hyperactivity: Part I. The Journal of pharmacology and experimental therapeutics. 2008;326(2):432–42. doi: 10.1124/jpet.108.139295. PubMed PMID: 18499743.

84. Nikolaev YA, Cox CD, Ridone P, Rohde PR, Cordero-Morales JF, Vasquez V, et al. Mammalian TRP ion channels are insensitive to membrane stretch. J Cell Sci. 2019;132(23). Epub 2019/11/15. doi: 10.1242/jcs.238360. PubMed PMID: 31722978; PubMed Central PMCID: PMCPMC6918743.

85. Sianati S, Schroeter L, Richardson J, Tay A, Lamande SR, Poole K. Modulating the Mechanical Activation of TRPV4 at the Cell-Substrate Interface. Front Bioeng Biotechnol. 2020;8:608951. Epub 20210118. doi: 10.3389/fbioe.2020.608951. PubMed PMID: 33537292; PubMed Central PMCID: PMCPMC7848117.

86. Nishida K, Nomura Y, Kawamori K, Moriyama Y, Nagasawa K. Expression profile of vesicular nucleotide transporter (VNUT, SLC17A9) in subpopulations of rat dorsal root ganglion neurons. Neurosci Lett. 2014;579:75-9. Epub 20140717. doi: 10.1016/j.neulet.2014.07.017. PubMed PMID: 25043192.

87. Woo SH, Lukacs V, de Nooij JC, Zaytseva D, Criddle CR, Francisco A, et al. Piezo2 is the principal mechanotransduction channel for proprioception. Nat Neurosci. 2015;18(12):1756–62. doi: 10.1038/nn.4162. PubMed PMID: 26551544; PubMed Central PMCID: PMCPMC4661126.

88. Mahmud AA, Nahid NA, Nassif C, Sayeed MS, Ahmed MU, Parveen M, et al. Loss of the proprioception and touch sensation channel PIEZO2 in siblings with a progressive form of contractures. Clin Genet. 2017;91(3):470–5. Epub 2016/09/09. doi: 10.1111/cge.12850. PubMed PMID: 27607563.

89. Chesler AT, Szczot M, Bharucha-Goebel D, Ceko M, Donkervoort S, Laubacher C, et al. The Role of PIEZO2 in Human Mechanosensation. N Engl J Med. 2016;375(14):1355–64. doi: 10.1056/NEJMoa1602812. PubMed PMID: 27653382.

